# Membrane association prevents premature degradation and mitigates inefficient biogenesis of suboptimal membrane proteins

**DOI:** 10.1101/2025.06.05.658084

**Authors:** Christian Diwo, Marta Alenquer, Colin Adrain, Maria João Amorim

## Abstract

Accurate membrane protein biogenesis is essential for cellular function, yet many proteins contain suboptimal targeting or insertion signals. Influenza A virus (IAV) faces similar constraints during infection but may have evolved strategies to enhance the biogenesis of its own membrane proteins. One such protein, the viroporin M2, contains functionally essential hydrophilic residues within its transmembrane domain, which should hinder efficient membrane insertion. We hypothesize that IAV has adapted to overcome these sequence-based limitations and ensure robust M2 biogenesis. Using a biotin pulse-labelling system in intact cells, we uncover the dynamics of M2 targeting and ER insertion. Attenuating the insertion rate by ablating a key insertion factor, the ER membrane protein complex (EMC), leads to M2 accumulating in the cytosol, but remaining partially insertion competent. We find that cytosolic stability of non-inserted M2 is crucial for efficient biogenesis under these conditions and identify amphipathic helix– mediated membrane association as the molecular mechanism that counteracts proteasomal degradation. We propose membrane association as a novel buffering mechanism that regulates membrane protein biogenesis by stabilizing pre-insertion intermediates.

**SIGNIFICANCE STATEMENT:** Membrane proteins failing to insert into the endoplasmic reticulum (ER) may be rapidly degraded to avoid protein aggregation in the cytosol. Our findings challenge this prevailing view. We show that the influenza A viroporin M2, despite possessing unfavourable features for membrane insertion, can increase biogenesis efficiency by associating with the membrane prior to insertion. Membrane association buffers the protein against quality control pathways and promotes successful biogenesis—even in the absence of a key insertion factor. These results reveal an alternative fate for non-inserted membrane proteins and uncover a viral strategy that redefines how cells manage unstable insertion intermediates.

## INTRODUCTION

Membrane proteins are synthesized by cytosolic ribosomes and must be efficiently inserted into their target membrane to avoid cytotoxic aggregation [1,2]. Non-inserted aggregation prone proteins like prion protein (PrP) may require fast removal but may not represent the broader class of membrane proteins. Recent studies have revealed a more nuanced picture: non-inserted or mislocalized membrane proteins are initially captured by cytosolic holdases such as SGTA [3], the BAG6 complex [4], and ubiquilins [5], which mask hydrophobic segments and temporarily stabilize these clients in the cytosol. From this buffered state, proteins may be inserted or targeted for degradation by the proteasome [6]. Thus, the cytosolic stability of non-inserted membrane proteins is not universally short-lived, but dynamically regulated and may serve as a critical checkpoint in membrane protein homeostasis. However, how protein features contribute to the stability of pre-insertion intermediates in the cytosol is poorly understood.

Fundamentally, the efficiency of membrane protein biogenesis is shaped by a protein’s sequence features and its interaction with available biogenesis machinery. Transmembrane domain (TMD) and signal peptide hydrophobicity [7,8], strongly influence membrane protein biogenesis efficiency [9–11] and influence targeting pathways and translocon selection [12–16].

The influenza A virus (IAV) proton channel M2 exemplifies this challenge. M2 is an essential component of the virion envelope, functioning in the uncoating step during viral entry and has been an effective antiviral drug target until the emergence of drug-resistant strains [17]. M2 is a 97-amino-acid, type III membrane protein with a single TMD that includes highly conserved hydrophilic residues [18]. These sequence features render M2 a suboptimal substrate for canonical targeting factors and insertases [19–23]. M2 sequence may have evolved other features to mitigate these suboptimal client features.

M2 encodes an amphipathic helix downstream of its transmembrane domain which has been extensively studied for its function during viral egress, enabling ESCRT-independent membrane scission [24], and is sufficient to localize cytosolic proteins to membranes [25]. Amphipathic helices are known to mediate membrane association [26,27], but have never been investigated as potential targeting features during membrane protein biogenesis. Using a biotin pulse-labelling assay to obtain kinetic measurements of glycosylation intermediates [28,29], we show that amphipathic-helix-mediated membrane tethering stabilizes non-inserted M2 in the cytosol and enhances its access to translocons, effectively buffering against biogenesis bottlenecks. These findings reveal a previously unappreciated intermediate state in membrane protein quality control to ensure robust insertion.

## RESULTS

### The loss of EMC affects M2 ER insertion rate

The EMC is the central insertase for transmembrane domains with short, translocated flanking domains at the ER [30]. In our accompanying study, the use of a glycosylation-competent M2 variant (M2.Y.F with a Leu2Asp mutation; Fig. 1A) was crucial in revealing the role of the EMC in M2 biogenesis and identifying compensatory insertion pathways. We hypothesized that the lack of EMC creates a membrane insertion bottleneck for M2. To test this, we sought to resolve the timing of M2 targeting and insertion, by complementing steady-state glycosylation analysis with a biotin pulse-labelling assay. M2.Y.F was N-terminally tagged with an AVI-tag (AVI.M2.Y.F, Fig. 1A), a short peptide sequence specifically biotinylated by birA ligase in the presence of biotin. This highly sensitive labelling approach minimally perturbs M2 biogenesis and preserves native cytosolic complexity (Fig. S1). Cytosolically expressed birA biotinylates AVI.M2.Y.F during the window when its N-terminus resides in the cytosol—beginning with emergence from the ribosomal exit tunnel and ending upon membrane translocation (Fig. 1B). M2’s translocation kinetics are observed via glycosylation. M2 in the cytosol is initially unglycosylated (-Y), will be glycosylated upon ER insertion (+Y) and further glycosylated when trafficked along the secretory route (++Y, Fig. 1B). Within each compartment, M2 could be subjected to degradation through different mechanisms (Fig. 1B) [31,32]

**Fig. 1.**
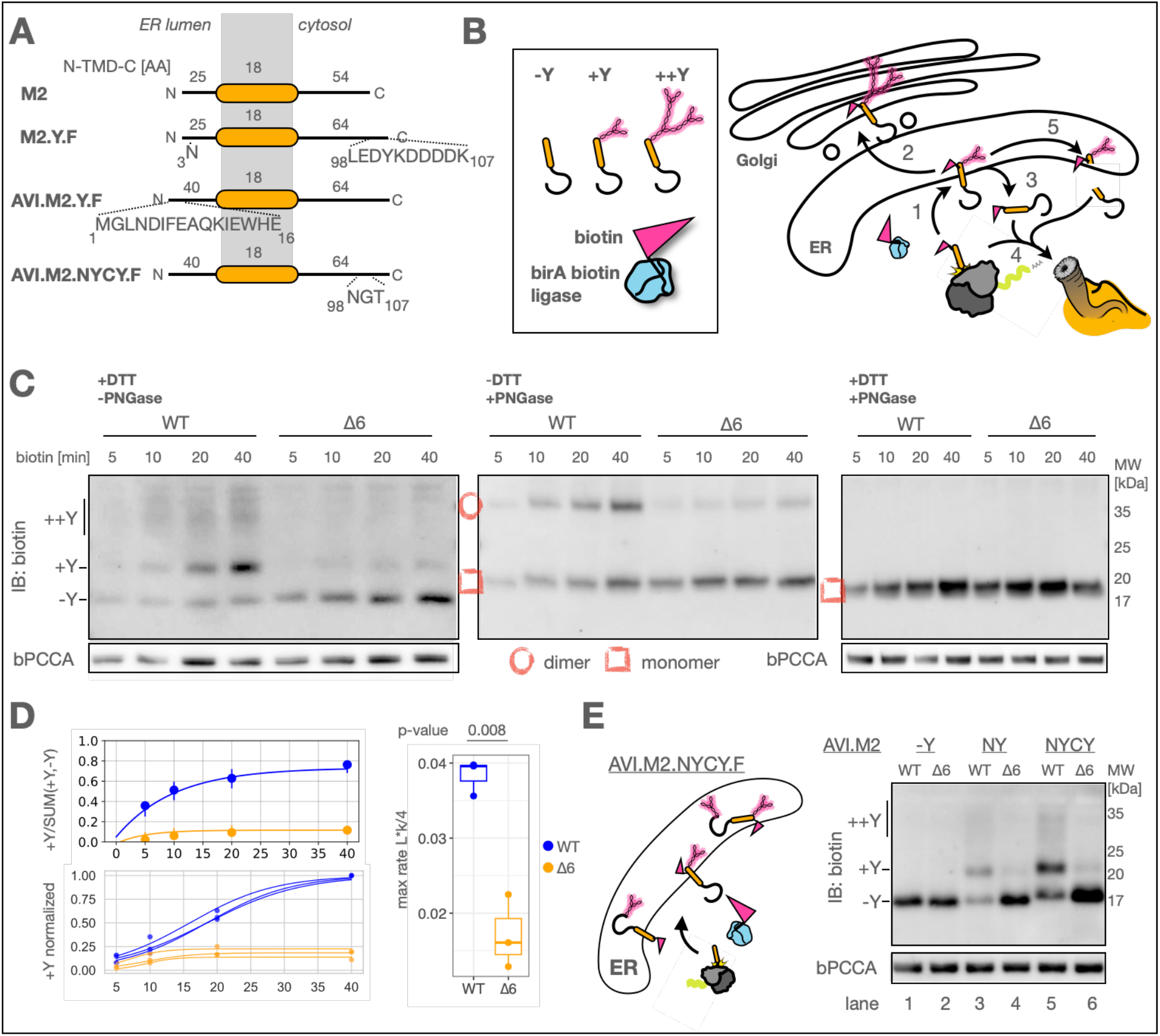
Pulsed biotinylation reveals M2’s ER membrane translocation dynamics. A) Scheme of M2 reporter constructs. M2 Leucine at position 3 is mutated to Asparagine and fused to a C-terminal Flag-tag (M2.Y.F). A 15 amino acid AVI-tag is fused to M2.Y.F’s N-terminus (AVI.M2.Y.F) which creates a highly specific biotinylation site for birA. An additional glycosylation site is inserted in M2.Y.F’s C-terminus (AVI.M2.NYCY.F). B) Scheme of possible reactions during M2 biogenesis acting on M2 with their results on glycosylation. 1) Membrane insertion 2) Trafficking 3) Retrotranslocation 4) Degradation 5) Cleavage. A549 WT and Δ6 cells were transfected with cytoBirA plus AVI.M2.Y.F for 24h, labelled with biotin for the indicated time intervals before harvesting on ice to quench the labelling reaction. The lysates were further treated either with DTT to dissociate M2 oligomers or PNGase to deglycosylate M2 and analyzed by western blot (N=3). D) Glycosylation efficiency (+Y/SUM(+Y,-Y)) and +Y levels were quantified by densitometry. An exponential model was fitted to the mean glycosylation efficiency of WT or Δ6. A logistic growth model was fitted to the mean +Y accumulation of WT or Δ6 of each repeat and data was normalized to the maximal estimated value and the maximal slope was determined as max accumulation rate. E) A549 WT and Δ6 cells were transfected with cytoBirA plus AVI.M2.-Y.F (no glycosylation site), or AVI.M2. Y.F (NY, regular N-terminal glycosylation site) or AVI.M2.NYCY.F (N- and C-terminal glycosylation site) for 24h, labelled with biotin for 5min before harvesting on ice to quench the labelling reaction. Lysates were analyzed by western blot (N=3).

A time course of pulsed biotinylation from 5 to 40 minutes in WT and EMC6 KO cells (Δ6) revealed a pronounced difference in the glycosylation dynamics. Whereas glycosylated species rapidly accumulated in WT cells, mostly unglycosylated species accumulated in Δ6 cells (+DTT -PNGase Fig. 1C). We quantified the glycosylation efficiency, which only reached 34% of the WT in absence of EMC (Fig. 1D). The maximal rate of M2 insertion reduced to 44% of the WT in Δ6 cells (Fig. 1D). In order to substantiate glycosylation as marker for ER insertion, we additionally measured disulphide-linked dimerization of M2, which occurs at positions C17 and C19 through the action of ER luminal isomerases [33]. The lack of EMC resulted in reduced dimerization (Fig. 1C -DTT +PNGase). Interestingly, our data showed that despite the highly reduced N-terminal translocation efficiency, unglycosylated M2 (-Y) accumulated at a rate that kept the total biotin labelled fraction close to equal between WT and Δ6 cells over the labelling period (Fig. 1C +DTT +PNGase), whereas the total M2 levels remain reduced (Fig. S1).

The lack of EMC has been reported to result in topologically inverted membrane insertion in vitro [13]. The accumulation of a non-glycosylated band may be consistent with a defect in topogenesis, thus we test if M2’s C-terminus may be translocated across the ER with a mutant that carries a glycosylation acceptor site at both N- and C-terminus (AVI.M2.NYCY.F, Fig. 1A). AVI.M2.NYCY.F is not glycosylated in a 10-minute biotin pulse in Δ6 cells (Fig. 1E lane 6), which excludes a defect in topogenesis. We conclude that a reduction in the rate of successful ER insertion reactions of M2, likely results in the accumulation of pre-insertion intermediates in the cytosol.

### M2’s amphipathic helix mediates membrane association and stabilizes non-inserted M2 in the cytosol

The accumulation of a stable unglycosylated M2 due to a defect in ER translocation is somewhat unexpected, as quality control mechanisms should rapidly degrade this fraction [1,2]. M2 sequences encodes an amphipathic helix directly following its transmembrane domain (Fig. 2A), which has been shown to target cytosolic red fluorescent protein to membranes [25]. We hypothesized that M2’s amphipathic helix may have a role in M2 biogenesis, potentially stabilizing non-inserted M2 in association with the cytoplasmic faces of the ER. We opted to construct a previously characterised amphipathic helix mutant, which maintains the secondary structure but abolishes membrane interaction by mutating 5 hydrophobic residues to alanine (AVI.M2-AHmut.Y.F, Fig. 1A) [34].

**Fig. 2.**
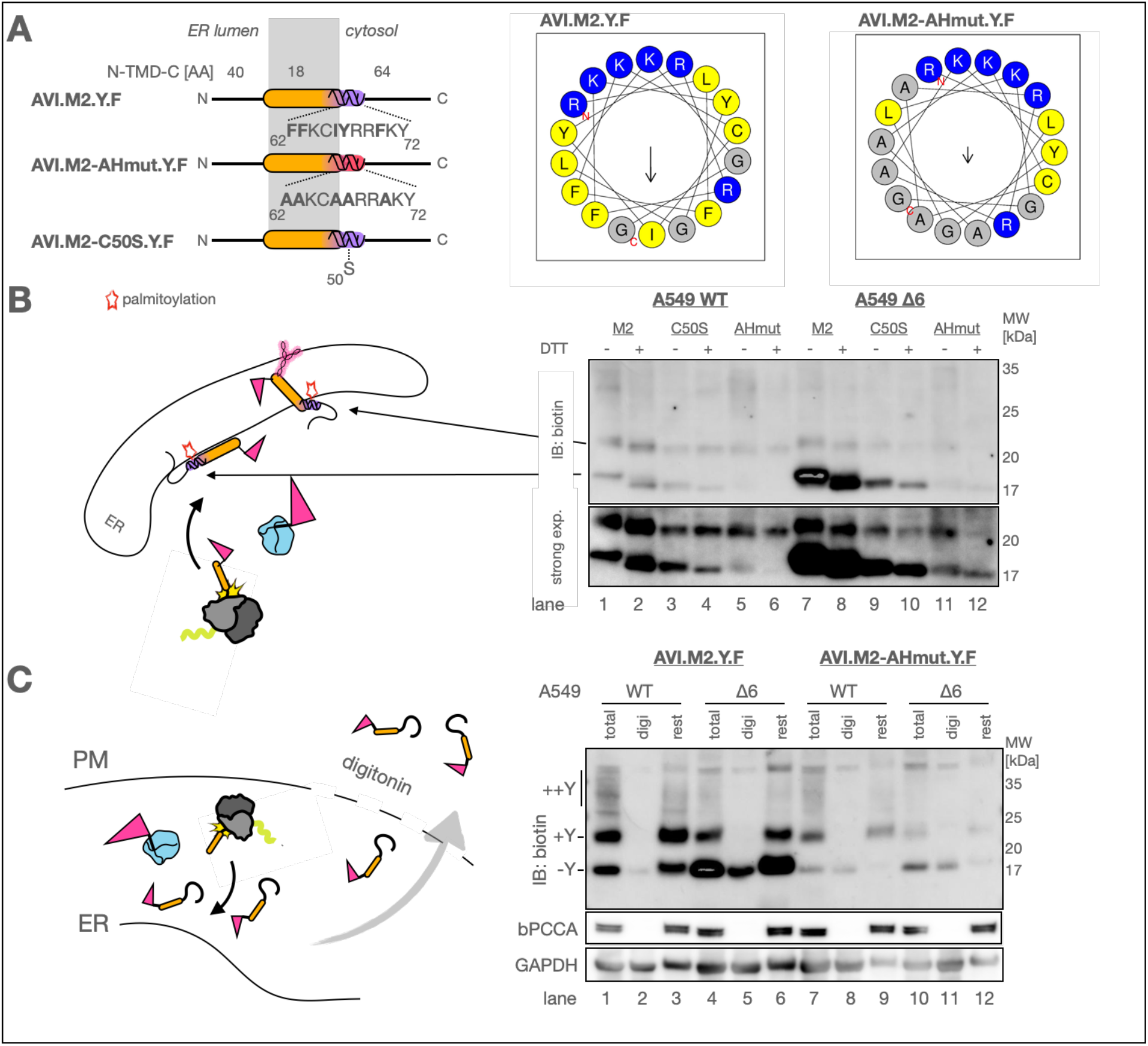
M2’s amphipathic helix mediated membrane association stabilizes non-inserted M2 in the cytosol. A) Sequence modifications of AVI.M2-AHmut and helical wheel representation created with heliquest [37]. B) A549 WT or Δ6 cells were transfected with cytoBirA plus AVI.M2.Y.F, or AVI.M2-C50S.Y.F or AVI.M2-AHmut.Y.F for 24h and labelled with biotin for 10min before harvesting on ice to quench the labelling reaction. Lysates were diluted in loading buffer with or without DTT and analyzed by western blot (N=3). C) A549 WT and Δ6 cells were transfected with cytoBirA plus AVI.M2.Y.F or AVI.M2-AHmut.Y.F for 24h, labelled with biotin for 40min before harvesting on ice with either 1% triton (total) or 0.02% digitonin. The digitonin solubilized cells were fractionated by centrifugation in supernatant (digi) and pellet (rest). Lysates were analyzed by western blot relative to the mitochondrial membrane protein PCCA (biotinylated bPCCA) and the cytosolic GAPDH (N=2).

First, we assessed if amphipathic helix mediates membrane association through monitoring palmitoylation, which occurs at a cysteine at position 50, located within its amphipathic helix [35]. Palmitoylation takes place at the cytosolic face of membranes and is catalysed by enzymes that localise to the ER, the Golgi, and the plasma membrane [36]. In agreement with membrane association, we detected palmitoylation as a shift in +Y and -Y fractions of AVI.M2.Y.F in both WT and EMC ablated conditions (Fig. 2B lane 1 and 7), compared to the DTT treated controls (Fig. 2b lane 2 and 8), or when the cysteine at position 50 is mutated to a serine (Fig. 2E lane 3,4 and 9,10). Strikingly, this band is almost entirely lost when M2’s amphipathic helix is disrupted (Fig. 2B lane 5,6 and 11,12), suggesting that M2’s amphipathic helix can stabilize -Y in association with the cytoplasmic faces of the ER.

To determine the subcellular localization of -Y, we selectively permeabilized the plasma membrane using digitonin and extracted the cytosolic contents. In WT cells, -Y was nearly undetectable in the cytosolic fraction (Fig. 2C, lane 2). In contrast, Δ6 cells showed partial detection of -Y in the cytosolic fraction (Fig. 2C, lane 5), while a strong -Y signal remained in the digitonin-resistant fraction in both WT and Δ6 cells (Fig. 2C, lanes 3 and 6). Notably, overall -Y protein levels were markedly reduced when expressing the AVI.M2-AHmut.Y.F construct under the same conditions as AVI.M2.Y.F. This was accompanied by a substantial decrease in both the cytosolic -Y (Fig. 2C, lanes 8 and 11) and the digitonin-resistant -Y (Fig. 2C, lanes 9 and 12).

These data suggest that amphipathic helix-mediated membrane association stabilizes fully translated M2 in association with the cytosolic membrane face of the ER, prior to post-translational membrane insertion. We test post-translational membrane insertion at the ER by inhibiting translation with a high dose of cycloheximide for 30 minutes, before pulse labelling cytosolic AVI.M2.Y.F with biotin. Indeed, we can observe glycosylation under these conditions in WT and Δ6 cells, confirming that at least a part of cytosolic -Y can insert post-translationally (Fig. S2).

### Cytosolic stability enhances biogenesis efficiency and delays proteasomal degradation

We hypothesize that an increased stability of pre-insertion intermediates in the cytosol may increase M2’s chance of ER insertion, and thus buffer against ER insertion bottlenecks. To assess how membrane association influences cytosolic stability and ER insertion rates, we pulse-labelled M2 and treated cells with either DMSO (mock), the proteasome inhibitor MG132, or the autophagy inhibitor bafilomycin A1 for 6 hours. The effectiveness of MG132 and bafilomycin A1 was confirmed by increased LC3 lipidation (LC3-II) compared to DMSO-treated controls (Fig. 3A). To rule out differences in transfection efficiency as a confounding factor, we confirmed comparable expression of co-transfected birA across all conditions (Fig. 3A).

**Fig. 3.**
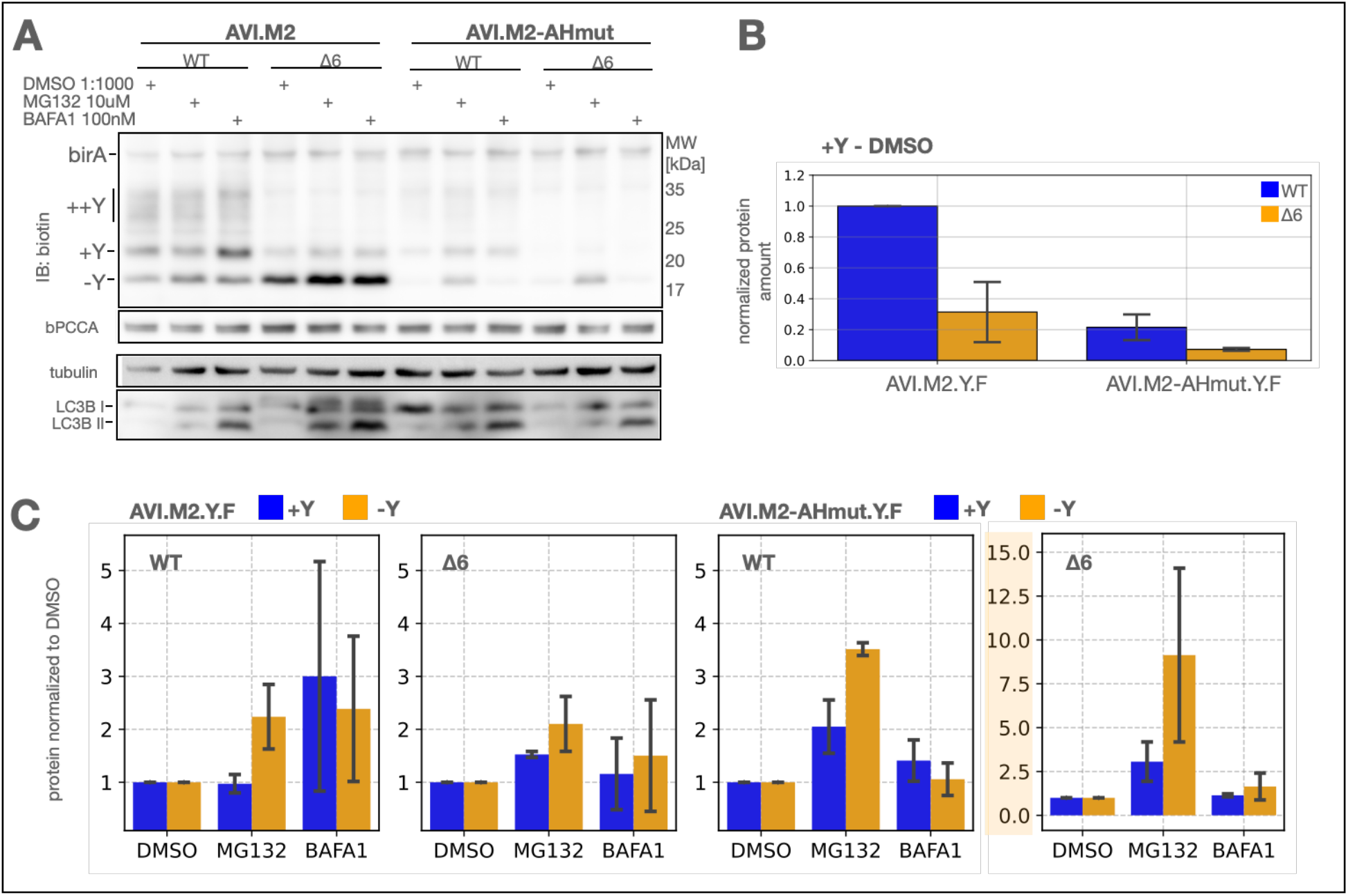
Impact of membrane association on ER insertion efficiency. A) A549 WT or Δ6 cells were transfected with cytoBirA plus AVI.M2.Y.F or AVI.M2-AHmut.Y.F for 20h and treated with either MG132, Bafilomycin A1 or the equivalent volume fraction of DMSO for 6h and labelled with biotin plus drugs for 20min before harvesting on ice to quench the labelling reaction (N=3). Lysates were analyzed by western blot. Bands were quantified by densitometry to assess ER insertion rate under control conditions B) or assess -Y stability C).

Consistent with previous experiments, we observed a substantial reduction in the insertion rate of +Y over 20 minutes in the absence of EMC (31% Δ6^M2^/WT^M2^, Fig. 3B). Notably, mutation of the amphipathic helix also significantly reduced M2 insertion (22% WT^AHmut^/WT^M2^, Fig. 3B). When both EMC was ablated and the amphipathic helix mutated, the insertion rate dropped even further (7% Δ6^AHmut^/WT^M2^, Fig. 3B). This multiplicative effect suggests that EMC and the amphipathic helix function independently, yet both are critical for efficient M2 insertion. We confirmed these findings using time-resolved biotin pulse-labelling, which showed a marked reduction in the maximal insertion rate of AVI.M2-AHmut.Y.F upon EMC ablation (19% Δ6/WT, Fig. S3). Together, these results highlight a strong contribution of membrane association— mediated by the amphipathic helix—to the biogenesis efficiency of M2, independent of translocation site availability.

If the cytosolic stabilization of a pre-insertion M2 aids in biogenesis efficiency, we should observe an increase in +Y levels through the inhibition of degradation. Proteasomal inhibition increased the -Y levels over DMSO treatment in all conditions (2x WT^M2^, 2x Δ6^M2^, 3.5x WT^AHmut^, 9x Δ6^AHmut^) and resulted in higher +Y levels relative to the DMSO treatment in all conditions except in WT cells expressing AVI.M2.Y.F (1x WT^M2^, 1.5x Δ6^M2^, 2x WT^AHmut^, 2.5x Δ6^AHmut^, Fig. 3A). Interestingly, lysosomal inhibition only increased AVI.M2.Y.F’s +Y levels in WT cells (3x WTM2, Fig. 3A), which is in agreement with previous research that links strong M2 expression with its lysosomal degradation [38]. We conclude that M2 can adhere to the cytosolic face of membranes via its amphipathic helix to escape proteasomal degradation, increasing its biogenesis efficiency.

### Lack of amphipathic helix renders infection more sensitive to EMC loss

M2 is highly expressed during PR8 infection, but only small amounts of M2 are needed at the plasma membrane to incorporate into virus particles [39,40]. If membrane association via M2’s amphipathic helix contributes to M2’s biogenesis efficiency, we should observe a defect in M2 plasma membrane accumulation when mutating the amphipathic helix. We created a mutant virus in the PR8 background carrying the same mutations in M2 that abolishes the hydrophobic features in the amphipathic helix (PR8-AHmut).

We infected WT and Δ6 cells, and measured HA and M2 accumulation at the plasma membrane together with viral titer at 8hpi. We assessed the magnitude of plasma membrane localization of these proteins in non-permeabilized cells in a flow cytometry assay and separated the infected cell population by modelling the expression profile with two skew-normal distributions, only considering the highly fluorescent population as infected cells (Fig. 4A). As expected, HA cell surface accumulation was not affected by EMC6 KO in both viruses. In contrast to the results we obtained in transfection, in infection M2-AHmut cell surface levels were not different from wild type M2 in WT cells. This suggests that IAV may regulate cellular processes that allow efficient biogenesis of M2 lacking the amphipathic helix in these cells. However, M2 accumulation at the plasma membrane was slightly more affected by EMC ablation when lacking the amphipathic helix (46% Δ6^PR8^/WT^PR8^ vs 28% Δ6^AHmut^/WT^AHmut^, Fig. 4B).

**Fig. 4.**
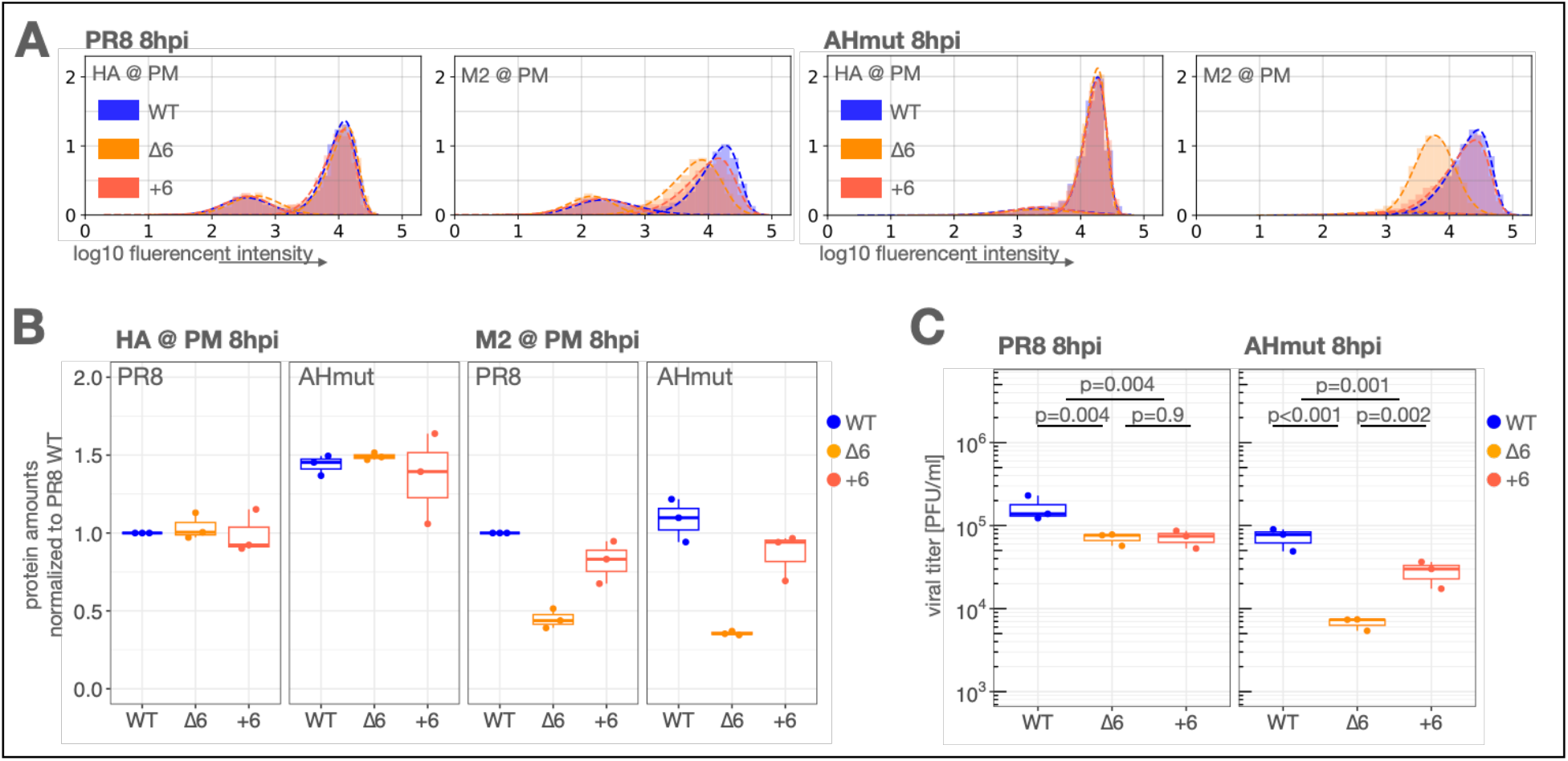
Role of M2 amphipathic helix in infection. A) HA and M2 levels at the surface of non-permeabilized infected cells (N=3). A549 WT, Δ6 or +6 cells were infected with PR8 or AHmut at MOI 3 for cell harvest at 8 hours post infection, and subsequent detection of viral HA and M2 by flow cytometry. The mean of an HA or M2 positive population of single cells was accurately estimated by fitting a skew normal distribution. B) Quantification of mean viral protein expression at the cell surface of A). C) Viral titers measured in supernatant in the same experiment described in A).

The loss of EMC function only modestly affected viral titer when comparing PR8 infected Δ6 to +6 cells (+6 = Δ6+EMC6 rescue cells). However, strikingly, the loss of EMC showed a significant impact on viral replication when M2’s amphipathic helix was mutated in PR8-AHmut replication (25% Δ6 PFU/+6 PFU, Fig. 4C). We observe similar effects in MDCK EMC6 KO cell lines (Fig. S5). These results demonstrate the functional relevance of M2’s amphipathic helix increasing M2’s biogenesis efficiency important to sustain sufficient ER insertion when insertion sites are scarce.

## DISCUSSION

Efficient membrane protein targeting and insertion are critical for cellular homeostasis, ensuring that thousands of simultaneously synthesized proteins are inserted into their correct organelles [41–43]. However, not all membrane proteins translocate efficiently, and weak sequence-encoded signals can result in toxic misfolded or non-inserted translation products [1,12,44,45]. Viruses, which rely on the host translocation machinery, must efficiently navigate these constraints to ensure proper membrane protein biogenesis of key viral proteins and virion assembly.

The relatively high efficiency of M2 biogenesis in absence of EMC led us to investigate potential redundancy in the membrane protein biogenesis and M2 features that would allow for partial compensation of EMC loss. Investigating M2’s ER translocation dynamics revealed that non-inserted M2 is initially stable in the cytosol rather than being rapidly degraded. Further analysis showed that cytosolic M2 can transiently associate with membranes via an amphipathic helix located near its transmembrane domain. Membrane association may hinder access of cytosolic ubiquitin ligases, prevent premature degradation, and thus increase the amount of insertion-competent substrate for translocon engagement. To the best of our knowledge, this has never been demonstrated.

The ability to buffer against limited translocation capacity may provide a significant advantage in viral infection. In our PR8 infection model, we show that M2 maintains sufficient plasma membrane levels to sustain viral replication in EMC-deficient cells. M2’s amphipathic helix is highly conserved across IAV strains, but certain mutations are common in viruses of specific hosts and serotypes (Fig. S4). Interestingly, the strength of its hydrophobic moment (strength of segregation of polar and apolar amino acids between the helix faces) varies between viruses of different host species, suggesting potential for adaptation (Fig. 5A). Other viral ion channels may exploit similar strategies. For example, HIV’s Vpu ion channel encodes two amphipathic helices in its C-terminus, suggesting that membrane association might be a conserved feature among viroporins [46].

**Fig. 5.**
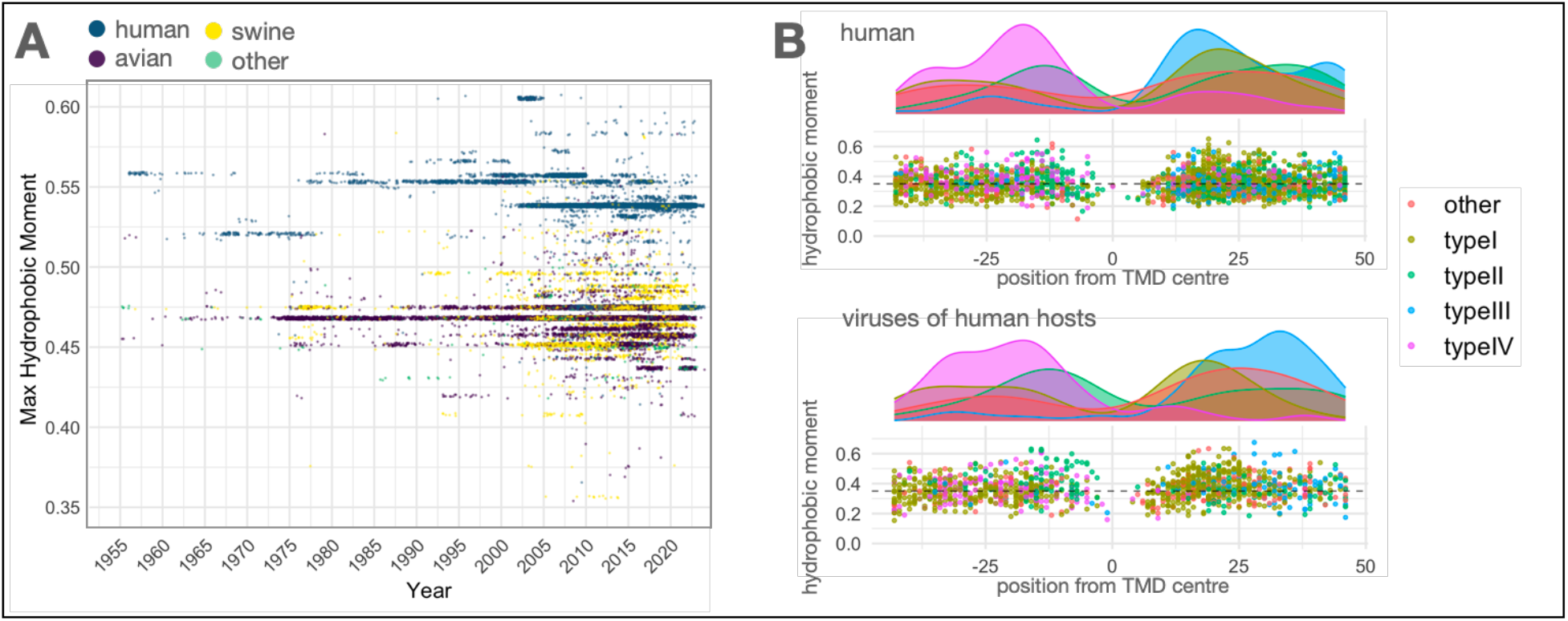
Sequence analysis of amphipathic features in human viral type III membrane proteins. A) 44621 M2 nucleotide sequences were aligned and translated computationally. Protein sequences with consensus 41WILDRLFFKCIYRRFKYGLKRG62 were analyzed for the maximal hydrophobic moment in a 12 amino acid window, assuming 3.6 residues per turn of a helix and applying the Eisenberg hydrophobicity scale [47]. Data was plotted by year of sampling and the organism category the sequence was sampled from indicated by color. A jitter around the y value was applied for better visualization. B) Sequences from the database described in the accompanying publication were analyzed in the same way as in A) along a 100 amino acid stretch centred around the TMD (typical TMD length being 20 amino acids). The dotted horizontal line at 0.35 is the mean hydrophobic moment for a large sample of random sequences.

Transmembrane domain-adjacent amphipathic helices may be a widespread feature. Computational analysis reveals that a significant portion of viral and human type III membrane proteins encode transmembrane domain-proximal amphipathic sequences (Fig. 5B). This suggests that translocation buffering via amphipathic helix-mediated membrane association could be a widespread adaptation among viruses and possibly some host proteins.

The M2 transmembrane domain must simultaneously function as a proton-conductive pore and provide a signal for membrane insertion—two roles that impose inherently conflicting sequence requirements. In particular, the hydrophilic residues critical for ion conductance can interfere with efficient membrane targeting. Our findings identify the M2 amphipathic helix as a novel structural element that compensates for suboptimal targeting signals, promoting membrane protein biogenesis when canonical insertion pathways are limited or saturated.

## LIMITATIONS OF THIS STUDY

Our study is solely based on the biogenesis dynamics of M2, and generalization of our results for other proteins needs experimental validation.

## MATERIALS AND METHODS

### Cell lines

A549 (ATCC CCL-185) and Madin-Darby Canine Kidney (MDCK.1, ATCC CRL-2935), were cultured in Dulbecco’s Modified Eagle Medium (DMEM, Thermo Fisher 21969-035) supplemented with 10% fetal bovine serum (FBS, Thermo Fisher 10500064), 2 mM L-glutamine (Thermo Fisher, 25030024), and 1% (v/v) penicillin-streptomycin (Biowest L0022-100) in a humidified incubator at 37°C and 5% CO2.

### Antibodies

anti-Flag-tag (Sigma-Aldrich F1804-200UG), anti-M2 14C2 (Abcam ab5416), anti-GAPDH (Sicgen, AB0049), streptavidin-HRP (Thermo Fisher S911), anti-LC3B (Cell Signalling 2775)

### Reagents

Avicel (Merck Supelco 11365-1KG), Formaldehyde (Acros 10231622), JetPrime (Polyplus 101000046), ECL Select (GE Healthcare RPN2235), BSA (Sigma-Aldrich A9418-100G), nitrocellulose membrane (GE Healthcare 10600003), FBS dialyzed (GE Healthcare HYCLSH30079.02), culture plates (Corning 734-1597), D-biotin (Supelco 000000000000047868), Phusion (NEB 174M0530L), Pfu Ultra (Agilent Technologies 600387), digitonin (Sigma-Aldrich D141), PMSF (Sigma-Aldrich 93482)

### Viruses

A/Puerto Rico/8/34 (PR8) virus were obtained via reverse genetics plasmid system (de Wit et al., 2007). 8 plasmids were transfected into 293T cells, and supernatants were used to expand in MDCK cells. The final viral stock solutions were obtained by using the supernatants from infected MDCK cells to infect embryonated chicken eggs. PR8-AHmut was obtained in the same way using PR8-Segment7-AHmut instead.

### Plasmids

Primers used for each construct are listed in Table S2. M2.Y.F was amplified with a forward primer adding the N-terminal AVI-tag and cloned into pcDNA3 to obtain AVI.M2.Y.F. The C50S mutation was introduced by quick change mutagenesis to obtain AVI.M2-C50S.Y.F. The glycosylation site in AVI.M2 was mutated back to the original leucine to obtain AVI.M2.-Y.F by quick change mutagenesis. AVI.M2.NYCY was obtained by introducing amino acids NGT at AVI.M2’s C-terminus before the Flag-tag by quick change mutagenesis. The mutations in M2’s amphipathic helix to obtain AVI.M2-AHmut.Y.F were introduced via overlap-PCR with primers containing the mutations and cloned into pcDNA3. The mutations in the IAV genomic segment 7,PR8-Segment7, were introduced in the same way to obtain PR8-Segment7-AHmut. birA was amplified from myc-birA (gift from Alfonso Bolado, Edinburgh) and tagged with an N-terminal V5-tag, and tagged with a C-terminal nuclear export signal (cytoBirA) and cloned into pcDNA3.

### Viral infection with samples for flow cytometry and plaque assay

Cells plated the previous day were washed in PBS and infected with viral solutions in serum free DMEM at MOI of 3. At the intended time points, cell supernatants were collected and frozen at - 80°C until use. For flow cytometry samples cells were detached in trypsin at 37°C for 7min and transferred to a 96 well plate with conical bottom to spin down at 300xg 4°C for 3min each spin. Cells were washed once in PBS + 1% FBS and once in PBS and fixed in PBS + 4% formaldehyde for 10min at room temperature. Cells were spun down at 500xg 4°C for 5min and washed twice in PBS and stored at 4°C for a maximum of 2 days. Cells were stained for 1h in primary antibody solution or PBS for unstained controls at room temperature. Cells were spun down and washed twice in PBS and stained with 1:1000 isotype specific secondary antibody solution Alexa 488 or Alexa 647 for 30min at room temperature. Cells were spun down and washed twice in PBS and measured at Fortessa X-20.

### Flow cytometry analysis

Data acquired on Fortessa X-20 was gated for cells and single cells using FlowJo. Single cells gated data for the two measured channels M2 and HA was exported and further processed in Python with custom scripts. Optimal parameters were fitted by maximum likelihood estimation for a mixture model of two skew-normal distributions using global optimization via differential evolution. Only the distributions with the larger mean at each time point were analyzed further. The data was processed in two different ways. First, the data for each channel was normalized to the 15h time point for each protein in the WT cells and the mean at each time point was compared between WT and Δ6 cells. Second, each repeat for each protein and WT or Δ6 condition was fit to a logistic growth model using a differential evolution algorithm and normalized to the parameter L of the WT condition for each protein and repeat. The data was fit again to obtain the final optimized parameters of the normalized data.

### Plaque assay

MDCK cells were seeded in 12 well plates to reach confluency the next day. Cells were washed in PBS and infected with 300ul of serially diluted supernatants in serum free DMEM. After 1h cells were washed in acid wash solution, washed in PBS and overlayed with serum free DMEM plus 0.14% BSA and AVICEL solution. Cells were incubated at 37°C + 5% CO^2^ for 30h.

### Pulsed biotinylation

A549 WT or Δ6 cells were seeded in 24 well plates (6×10^4 cells/well) and transfected 16h later with cytoBirA plasmid and AVI.M2.Y.F or AVI.M2-AHmut.Y.F plasmid (each 150ng/well) with Jetprime reagent and incubated for 20h in pre-labelling media (DMEM +2mM GLUT +1%P/S +10%FBS dialysed). Medium was replaced with labelling media +100nM biotin for the time indicated in the experiment, quickly placed on ice and washed 2x with cold PBS to quench the biotinylation reaction. From here on all steps were performed on ice or 4°C. Cells were lysed in 70ul TX100 lysis buffer (50mM Tris-HCl pH 7.4, 150mM NaCl, 1% (v/v) Triton X-100, 5mM EDTA and protease inhibitors) on ice, scraped and transferred to 1.5ml test tubes and spun at 1000xg and 4°C. The supernatant was transferred to new tubes, mixed with 6x LDS sample buffer (550 mM Tris-HCl pH 7.4, 200 mM LDS, 0.13 mM EDTA, 250 mM DTT, 15% v/v glycerol, 0.025% SERVA Blue G250 and 0.025% Phenol Red) and transferred from ice to a 95°C hot plate for 3min. Samples were stored at -20°C until analysis by western blot.

10% Acrylamide gels containing SDS and sucrose were loaded with samples and run in MOPS running buffer (50 mM MOPS, 50 mM TrisBase, 3.5 mM SDS, and 0.8mM EDTA) at 150V for 50min. Gel was transferred in Biorad transfer system onto nitrocellulose membranes, blocked in 5% milk in PBS + 0.1% Triton X-100 and incubated in primary antibody overnight at 4°C or 1:10000 streptavidin-HRP for 1h at room temperature. 1:10000 secondary antibody incubation was 1h at room temperature. Chemiluminescent detection was performed on Amersham AI600. Western blots were quantified by densitometry using FIJI ImageJ.

### Cytosol extraction with digitonin

Cells were treated following pulsed biotinylation for 40min after quenching the reaction on ice. From here on all steps were performed on ice or 4°C. For the total fraction, cells were lysed in 70ul TX100 lysis buffer on ice, scraped and transferred to 1.5ml test tubes. To obtain the digi and rest factions, 50ul PBS + 0.02% digitonin + 1mM PMSF was added to each well and incubated on ice for 20min. The supernatant was collected in a 1.5ml tube, and cells left in the well were lysed in 50ul TX100 lysis buffer and transferred to a 1.5ml tube. Total, digi and rest fractions were spun down 10min at max speed. The supernatants of all fractions were transferred to new tubes, mixed with 6x LDS buffer and transferred from ice to a 95°C hot plate for 3min. Samples were stored at -20°C until analysis by western blot.

### Computational analysis of amphipathic properties

44621 IAV segment 7 nucleotide sequences were obtained from NCBI (downloaded), aligned and translated. Residues 41-62 of the M2 open reading frame were analyzed. The single pass protein database analyzed in our accompanying publication was analyzed for potential transmembrane domain proximal amphipathic helices along a 100 amino acid sequence centered around the transmembrane domain center assessed by the ΔGapp minimum. Hydrophobicity was assessed through the Eisenberg scale [47] assuming 3.6 residues per turn of a helix.

### Statistical analysis

The statistical analysis used is detailed in each figure legend.

## ACKNOWLEDGEMENTS

This project has received funding from the European Research Council (ERC) under the European Union’s Horizon 2020 research and innovation programme (grant agreement No 101001521) to M.J.A., ‘la Caixa’ Foundation project grant CF/PR/HR17/52150018 to C.A. and Fundação para a Ciência e a Tecnologia 2020.10234.BD to C.D‥ Bafilomycin A1 and MG132 were a kind gift from Moritz Treeck. We thank Ignacio González Bravo for critical discussions. We thank Tiago Paixão for his insights into the data analysis and guidance throughout the project. Author contributions: C.D., C.A., and M.J.A. conceptualized the research; C.D. and M.A. performed research; C.D., and M.A. contributed new reagents; C.D., M.A., and M.J.A analyzed data; C.D. wrote the manuscript; and C.D., C.A., and M.J.A. revised the manuscript. Competing interests: The authors declare no competing interest.

