## Supplementary material for "Membrane association prevents premature degradation and mitigates inefficient biogenesis of suboptimal membrane proteins": SI

SUPPLEMENTAL INFORMATION

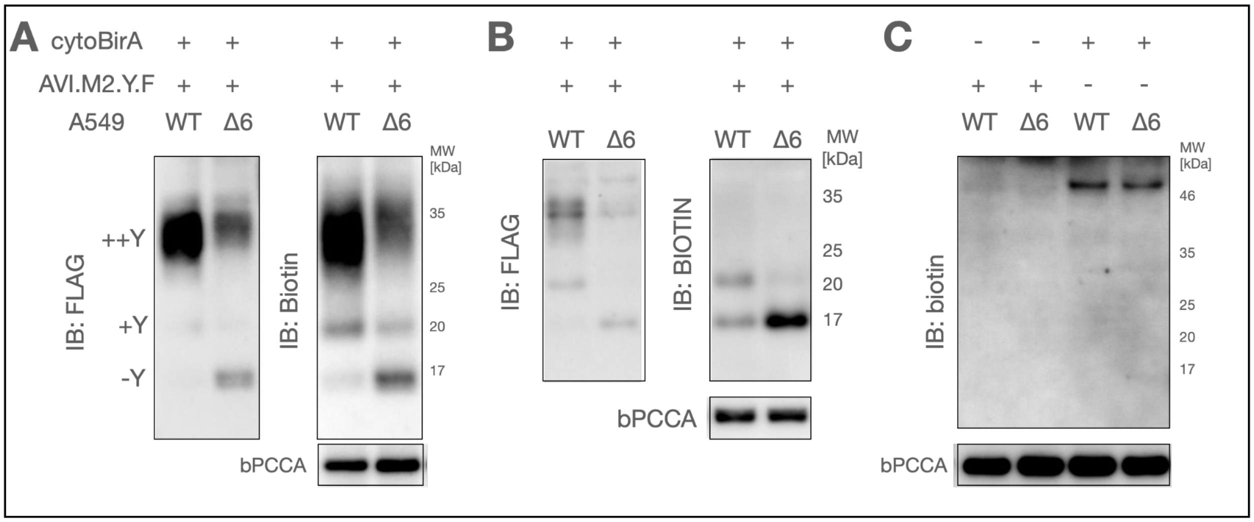


Fig. S1. AVI.M2.Y.F specific biotinylation by birA. A549 WT or Δ6 cells are transfected with cytoBirA plus AVI.M2.Y.F for 24h and labelled for 24h in A) or 5min in B) with biotin. Cells were lysed on ice to quench the reaction and analysed by western blot. Blots were first stained with anti-FLAG, imaged, stripped and stained again with streptavidin-HRP and imaged again. C) Cells were transfected with only AVI.M2.Y.F or only cytoBirA, labelled for 24h with biotin and analysed by western blot.

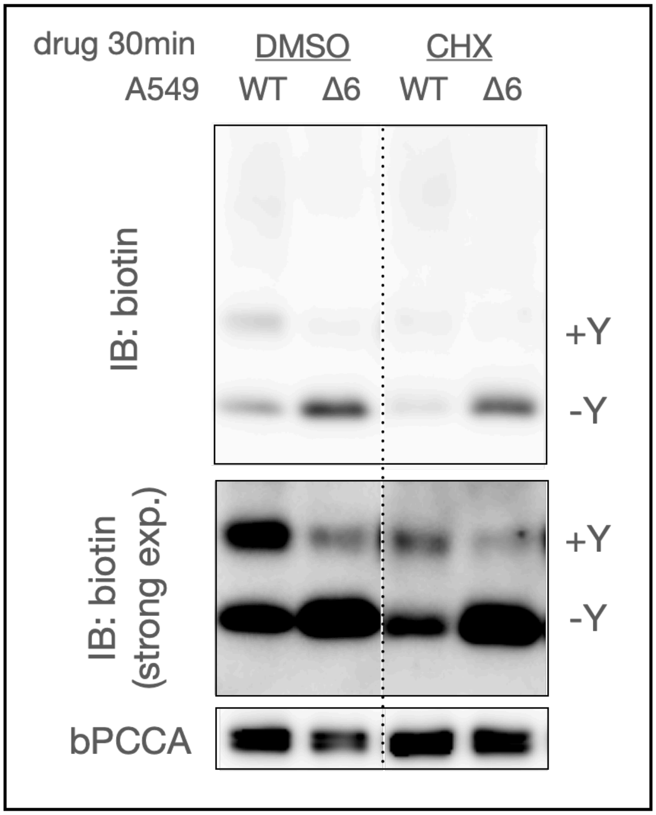


Fig. S2. Post-translational insertion of M2. A549 WT or Δ6 cells were transfected with cytoBirA plus AVI.M2 for 20h and treated with either 50ug/ml CHX, or the equivalent volume fraction of DMSO for 30 min, and labelled with biotin plus drugs for 20min before harvesting on ice to quench the labelling reaction. Lysates were analysed by western blot (N=2).

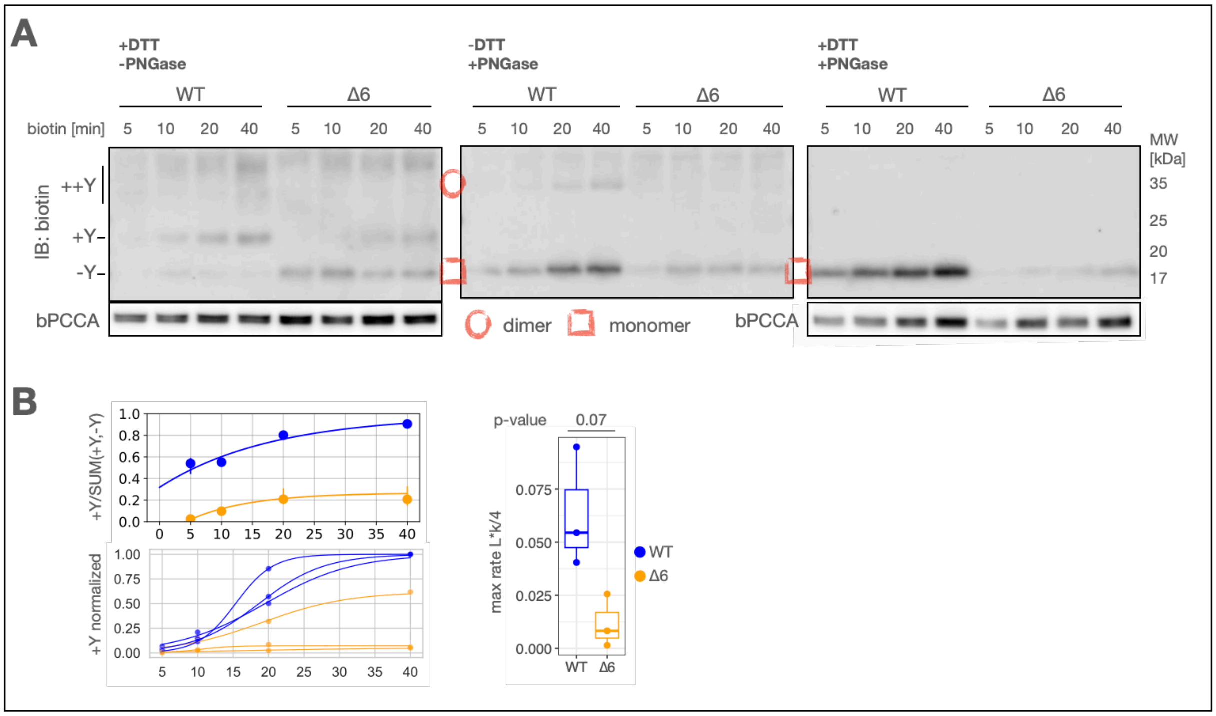


Fig. S3. ER biogenesis dynamics of AVI.M2-Ahmut.Y.F. A) A549 WT and Δ6 cells were transfected with cytoBirA plus AVI.M2-AHmut.Y.F for 24, labelled with biotin for the indicated time intervals before harvesting on ice to quench the labelling reaction. The lysates were treated ±DTT and ±PNGase and analysed by western blot (N=3). B) Glycosylation efficiency (+Y/SUM(+Y,-Y)) and +Y levels were quantified by densitometry. An exponential model was fitted to the mean glycosylation efficiency of WT or Δ6. A logistic growth model was fitted to the mean +Y accumulation of WT or Δ6 of each repeat and data was normalized to the maximal estimated value and the maximal slope was determined as max accumulation rate.

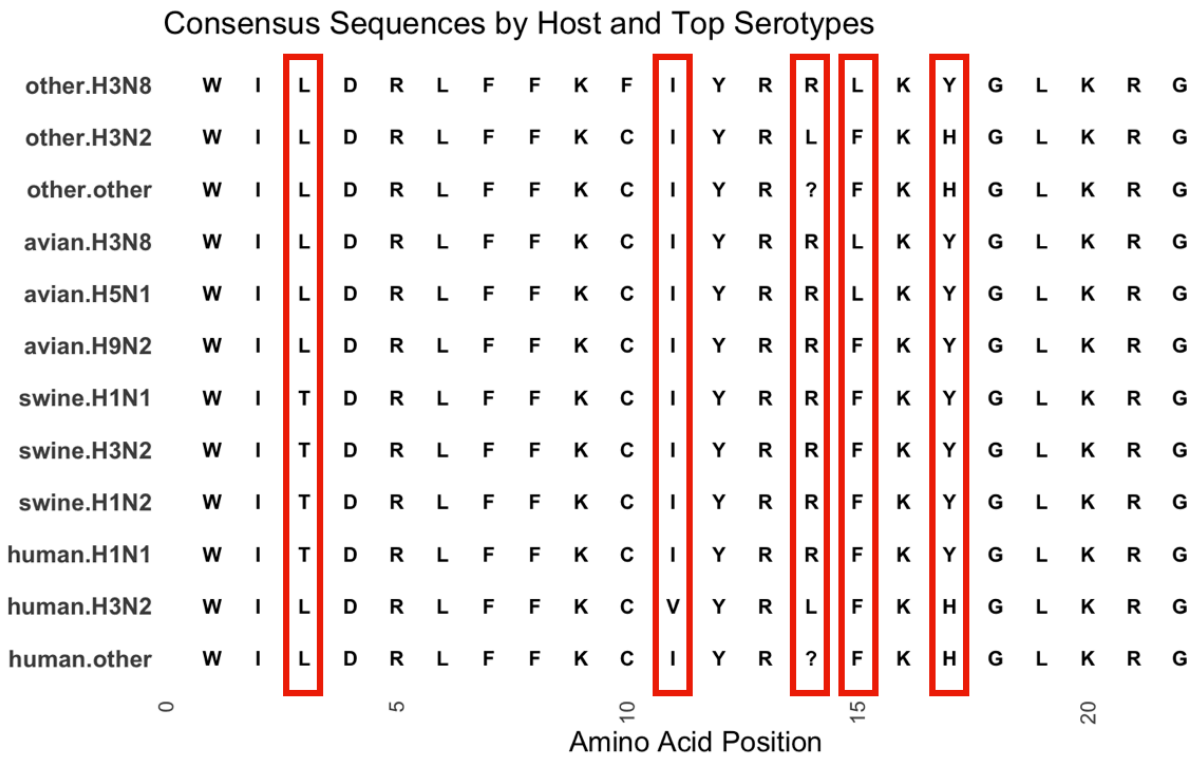


Fig. S4. Mutations underlying differences in hydrophobic moments of IAV amphipathic helix. Consensus sequences of amino acids position 41-62 in M2 for the most prevalent serotypes within each host category. Same set of sequences from Fig. 5A.

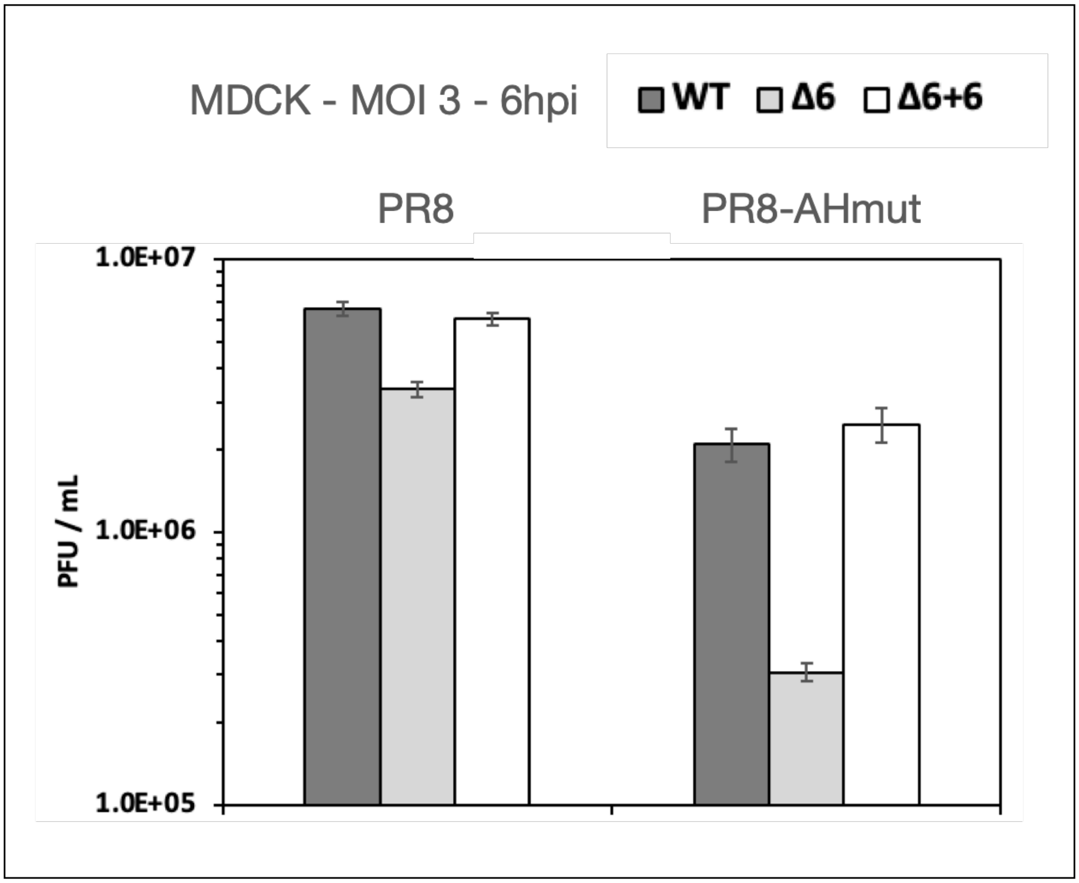


Fig. S5. Viral infection defect due to lack of EMC6 in MDCK. MDCK WT, △6 or +6 cells were infected with PR8 or PR8-AHmut at MOI 3 for 6h and supernatants were collected and assessed by plaque assay (N=1).
